# ComEB protein is dispensable for transformation but must be translated for optimal synthesis of ComEC

**DOI:** 10.1101/2020.12.30.424838

**Authors:** Micaela De Santis, Jeanette Hahn, David Dubnau

**Affiliations:** Public Health Research Institute Center, New Jersey Medical School, Rutgers University, Newark, NJ, USA

**Keywords:** transformation, translational coupling, nucleotide scavenging, ComEB, ComEC, post transcriptional regulation

## Abstract

We show that the ComEB protein is not required for transformation in *Bacillus subtilis,* despite its expression from within the *comE* operon under competence control. We show further that the synthesis of the putative channel protein ComEC is translationally coupled to the upstream *comEB* open reading frame, so that translation of *comEB* and a suboptimal ribosomal binding site embedded in its sequence are needed for proper *comEC* expression. Translational coupling appears to be a common mechanism in three major competence operons for the adjustment of protein amounts independent of transcriptional control, probably ensuring the correct stoichiometries for assembly of the transformation machinery. *comEB* and *comFC* respectively encode cytidine deaminase and a protein resembling type 1 phosphoribosyl transferases and we speculate that nucleotide scavenging proteins are produced under competence control for efficient reutilization of the products of degradation of the non-transforming strand during DNA uptake.

## Introduction

Bacterial transformation is an important mode of horizontal gene transfer and a complete understanding of its mechanism requires the identification of all the genes and proteins required for the binding, uptake and integration of transforming DNA. Although most of the essential players have been identified (Blokesch, 2016, Dubnau & Blokesch, 2019, Maier, 2020), uncertainty remains in the case of ComEB.

The conserved ComEA and ComEC proteins, are needed for transformation in nearly all bacteria, and at least in the firmicutes are encoded in the *comE* operon, which also encodes ComEB. ComEA is a DNA binding protein that participates in the uptake of transforming DNA to the periplasm (Provvedi & Dubnau, 1999, Gangel *et al*., 2014, Matthey & Blokesch, 2016, Schwarzenlander *et al*., 2009), while ComEC is a large, multipass membrane protein that most likely forms a channel for DNA transport to the cytoplasm (Draskovic & Dubnau, 2005). In *Bacillus subtilis* and in many other firmicutes, the *comE* operon also encodes ComEB, a dCMP deaminase (Burghard-Schrod *et al*., 2020, Oehlenschlaeger *et al*., 2015), while in other bacteria only *comEA* and *comEC* are present. It has been suggested that ComEB is dispensable for the transformation of *B. subtilis,* because a short in-frame deletion *(comEB^Δ84^),* removing 28 out of 189 codons from the 5’ end of the gene, had no effect on transformability (Inamine & Dubnau, 1995). Although this conclusion is consistent with the absence of *comEB* in some transformable bacteria that encode ComEA and ComEC, it remained possible that the major portion of ComEA still encoded by *comEB^Δ84^* was sufficient for normal transformability. A recent report that ComEB plays an important role in transformation (Burghard-Schrod *et al*., 2020) suggests that this may indeed be the case and caused us to re-examine the role of this protein.

We show here that the ComEB protein is completely dispensable for transformation. We also show that the *comEB* and *comEC* open reading frames are translationally coupled, explaining why deletion of *comEB* causes a loss of transformability. Finally, we speculate as to the reason for coupling and for why ComEB, a dCMP deaminase (Burghard-Schrod *et al*., 2020, Oehlenschlaeger *et al*., 2015), is expressed by cells that are competent for transformation.

## Results

In *B. subtilis, comEB* terminates in a pair of UGA stop codons that are immediately followed by the start of *comEC* (**UGA UGA** *AUG),* where the stop codons of *comEB* are in boldface and the start of *comEC* is italicized (Fig. 1A). This arrangement suggests that a ribosomal binding site (RBS) for *comEC* may be embedded in the coding sequence of *comEB* and even that the two genes may be translationally coupled, so that the translation of the *comEB* coding sequence is required for translation of *comEC* as suggested previously (Inamine & Dubnau, 1995). The transformation deficient *ΔcomEB::tet* construct used by Burghard-Schrod et al (2020) removed the entire *comEB* reading frame including any sequences embedded in *comEB* that might contain an RBS for *comEC* translation. To explore the role of *comEB* we utilized a number of constructs, all replacing the native locus in single copy, under normal competence control (Fig. 2). Previous work has shown that the *comE* operon is transcribed from a promoter upstream of *comEA*, with a likely additional minor promoter located between *comEA* and *comEB* (Hahn *et al*., 1993) (Fig. 2). Thus, the deletion of sequences within *comEB* is *a priori* unlikely to interfere with the transcription of either *comEA* or *comEC.*

**Figure 1.**
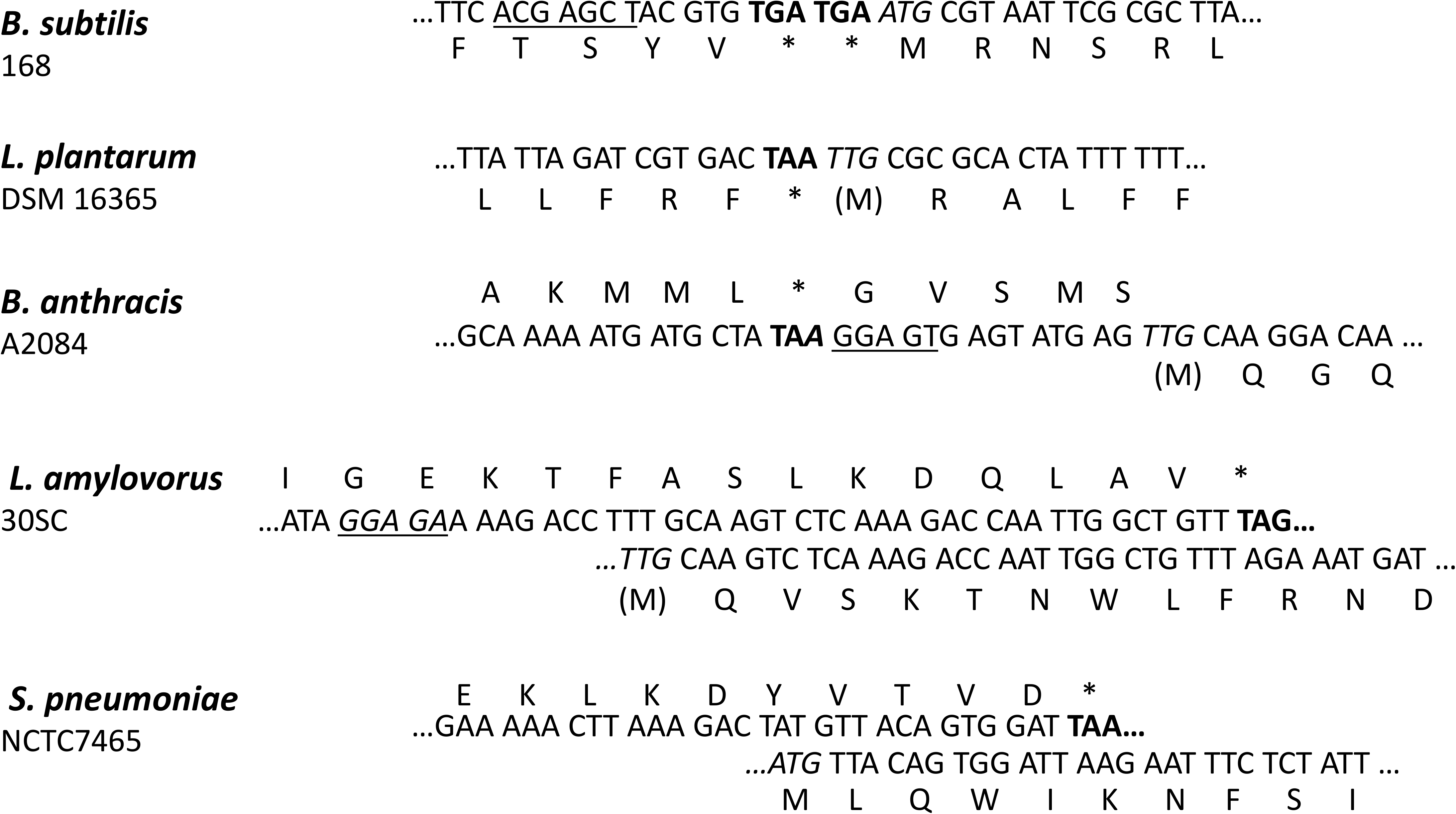
Arrangements of *comEB* and *comEC* in various firmicutes, identified with their strain numbers. In each case the last few codons of *comEB* and the first few codons of *comEC* are displayed together with their translated products. The *comEB* stop codons are in boldface, the *comEC* start codons are in italics and the proposed Shine-Dalgarno motif for *B. subtilis* is underlined. In (B), (C) and (D), the initial amino acid is shown in parentheses, because the TTG start is nominally translated as leucine. Putative Shine-Dalgarno sequences are underlined, when present.

The *ΔcomEB::ery* construct (Fig. 2) replaces nearly the entire coding sequence of *comEB* with an erythromycin resistance cassette, without perturbing the coding sequences of *comEA* and *comEC* (Koo *et al*., 2017). The construct retains the start codon of *comEB* and leaves 21 base pairs from the end of the *comEB* open reading frame. Although *ery* is oriented in the same direction as *comEB,* a stop codon is present at the end of the cassette, which will cause a translating ribosome to detach before reaching the final 21 residues of *comEB.* In several experiments with this construct, a low level of transformation was measured, about 40-fold lower than that of the wild-type strain, but still well above the transformation frequency of a *ΔcomEC* mutant, which is completely deficient (Table 1). This result confirms the finding of Burghard-Schrod *et al* (2020), which led to the conclusion that the ComEB protein is needed for transformation, admitting two possible interpretations. Either ComEB protein is needed for transformation or the deletion has removed sequences that are essential for the expression of *comEC.* Note that in the *ΔcomEB::ery* construct, the promoter associated with the resistance cassette is expected to drive downstream genes (Koo *et al*., 2017), so that the cassette should not be transcriptionally polar on *comEC*. To test whether the *ΔcomEB::ery* mutation reduced the translation of *comEC,* as suggested by the contiguity of their coding sequences, we used Western blotting with an antiserum against a twin-strep tag (TST) fused to the C-terminus of ComEC. Because ComEC is weakly expressed, we used a host carrying a *Pmtl-(comK comS)* construct, to increase expression of competence proteins (Rahmer *et al*., 2015). Fig. 3 shows that the *ΔcomEB::ery* strain produced no detectable ComEC, consistent with the idea that removal of *comEB* sequences prevents the full expression of *comEC.* Some undetected ComEC must be present because the transformation frequency is not zero (Table 1).

**Figure 2.**
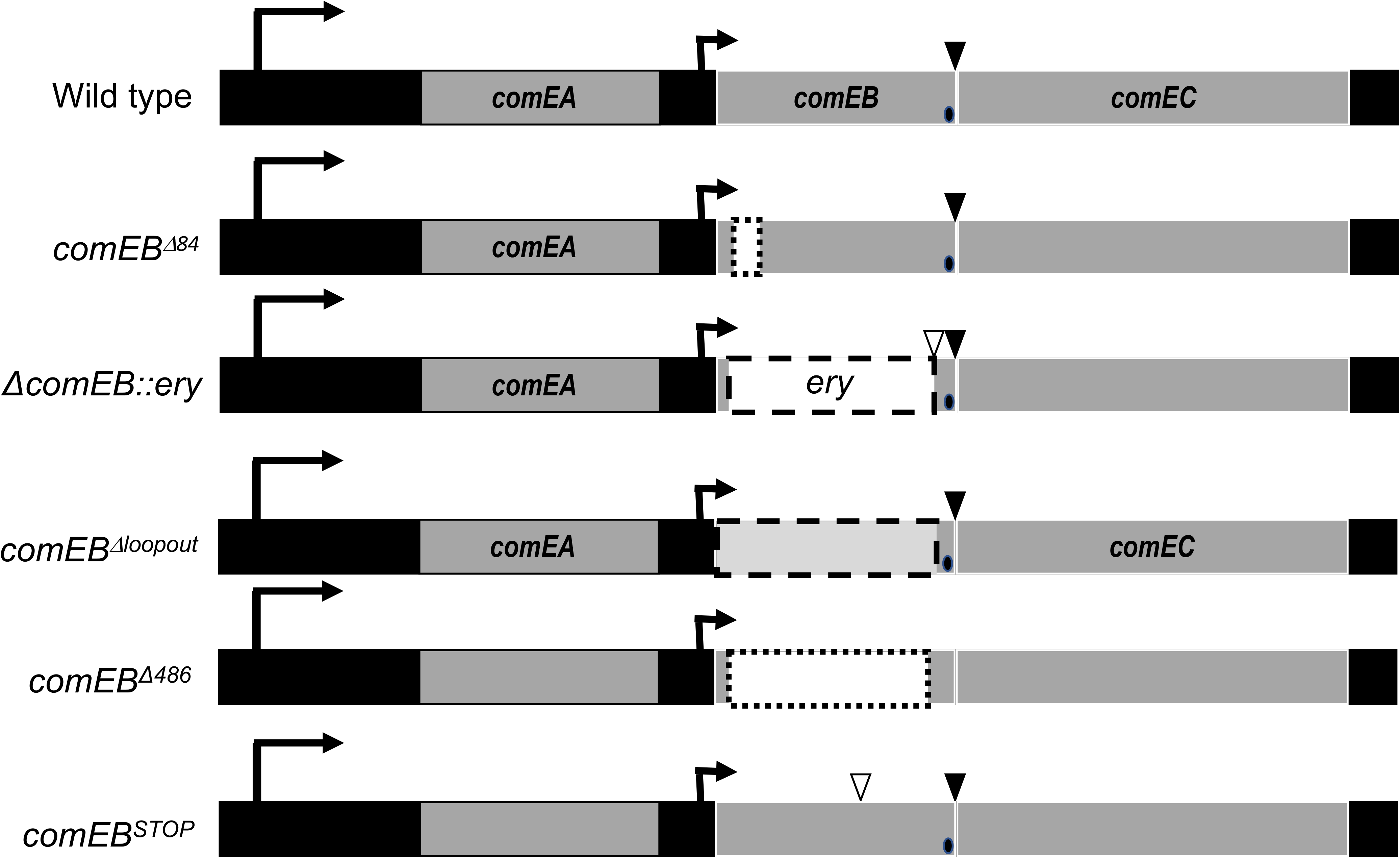
*comEB* constructs referred to in the text. The top diagram shows the wild-type *B. subtilis comE* operon. Dotted line boxes show deleted sequences, and the dashed boxes show replacements of *comEB* by the *ery* cassette and the scar left by the *comEB^Δloopout^* construct. Arrows show the locations of the *comE* major and minor promoters (Hahn *et al*., 1993). The *comEB^Δ84^, comEB^looput^* and *comEB^Δ486^* constructs leave sequences inframe with the start of *comEB* and contain the final base pairs of *comEB,* including the putative *comEC* RBS, depicted by black ellipses. The *comEB^Δloopout^* construct was derived from *ΔcomEB::ery* by removal of the *ery* cassette. This event leaves behind a scar of 150 in-frame residues, indicated by the light gray box. The open and closed triangles show introduced and naturally occurring stop codons, respectively. The reading frames and deletions are not drawn to scale. All of the mutant constructs replaced the wild-type sequence in single copy at the native locus, under normal competence control.

**Fig. 3.**
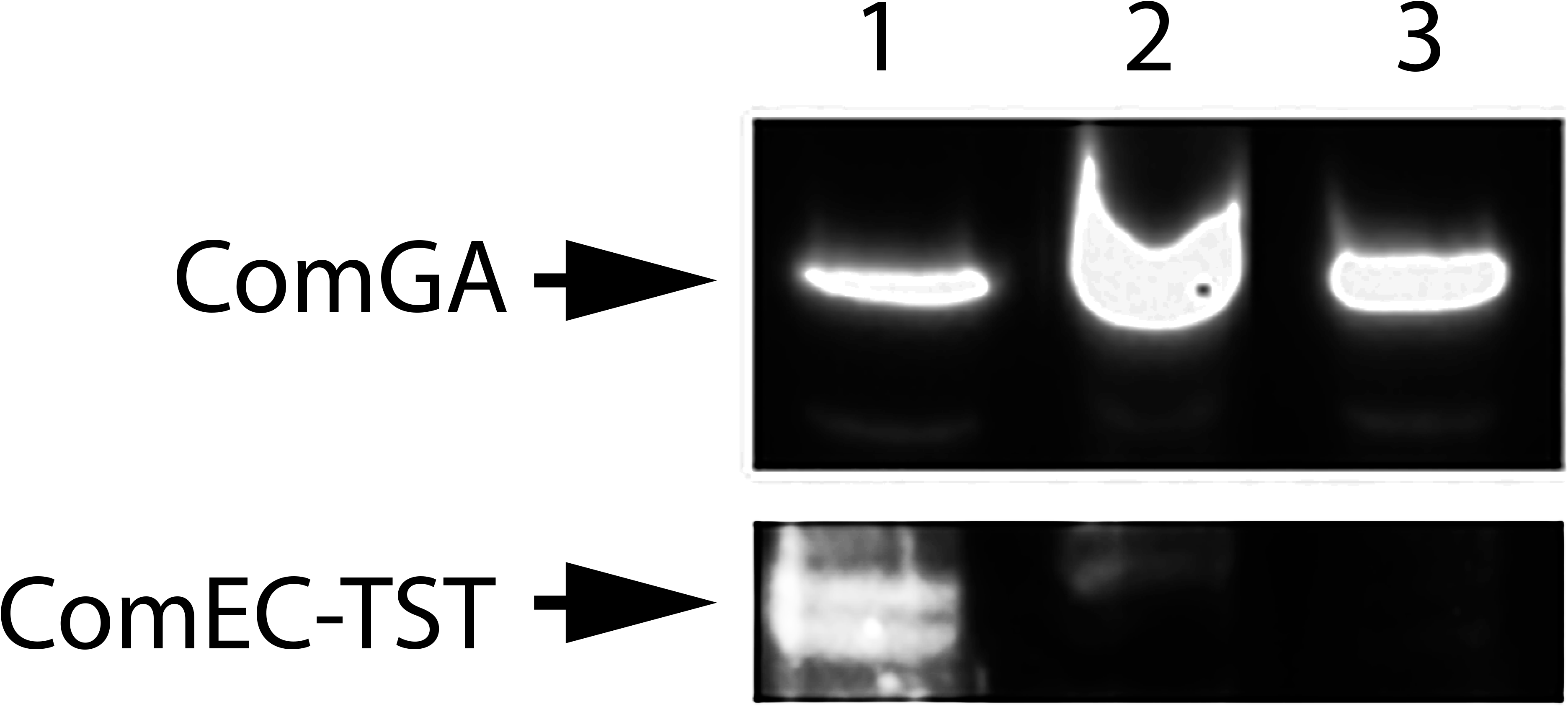
A *ΔcomEB::ery* strain (BD8953) produces less ComEC-TST than its wild-type parent (BD8931). Cultures were grown in LB and induced by growth in the presence of mannitol. Membrane fractions were isolated and Western blotting was carried out with an antiserum against the TST of both the wild-type (lane 1) and the *comEB* mutant (lane 2) strains. A *Pmtl-(comK comS)* strain lacking the TST fusion was included to show the specificity of the antiserum (lane 3). Antiserum raised against the membrane protein ComGA was used as a loading control.

**Table 1.** Transformation of *comE* mutant constructs

| ***comE* genotype** | **Transformation frequency ± SD<sup>a</sup>** |
| --- | --- |
| Wild-type (IS75) | 1.00 |
| $\Delta comEB::ery$ | $0.026 \pm 0.004$ |
| *comEB*<sup>*Δloopout*</sup> | $0.27 \pm 0.14$ |
| *comEB*<sup>*Δ486*</sup> | $3.0 \pm 1.4$ |
| *comEB*<sup>*STOP*</sup> | $0.0085 \pm 0.072$ |
| $\Delta comEC::ery$ | $<10^{-5}$, <sup>b</sup> |

<sup>a</sup>For each determination, the number of colony forming units ml<sup>-1</sup> of Leu<sup>+</sup> transformants was divided by the total colony forming units ml<sup>-1</sup> determined using LB agar. The values for each strain were normalized to that of the wild type strain in the same experiment, which exhibited an average frequency of $6.9 \times 10^{-4} \pm 2.9 \times 10^{-4}$ across all the experiments. Each determination was performed at least three times and average normalized values are shown together with standard deviations.
<sup>b</sup> No transformant colonies were detected in 100 μl of culture after transformation.

To further between the two interpretations described above, we used the *cre*-expressing plasmid pDR244 (Koo *et al*., 2017), to generate the *comEB^Δloopout^* construct that removes the *ery* cassette together with its stop codon (Fig. 2). The resulting strain carries the start codon of *comEB,* a 150 base pair scar from excision of the cassette and the last 21 base pairs of *comEB,* which are placed in-frame with the *comEB* start codon to produce a theoretical protein of 58 residues. This strain restored transformation to a level an order of magnitude higher than *ΔcomEB::ery,* although about 4-fold lower than the wild-type strain (Table 1). As a further demonstration that ComEB is not needed for transformation, we constructed *comEB^Δ486^,* an in-frame deletion construct that leaves the first ten codons of *comEB* and 54 residues at the downstream end of the gene, thus encoding an in-frame theoretical peptide of 28 residues in place of the 189 residue native ComEB (Fig. 2). As shown in Table 1, the *comEB^Δ486^* strain, like *comEB^Δloopout^,* is transformable, and in fact consistently exhibits a higher transformation frequency than the wild type strain. The phenotypes of the *comEB^Δloopout^* and *comEB^Δ486^* strains, together with the previous report that *comEB^Δ84^* is fully transformable (Inamine & Dubnau, 1995), show conclusively that ComEB is not needed for transformation.

As noted above, the *ΔcomEB::ery* construct inserts the *ery* cassette with its stop codon, so that the final 21 base pairs at the downstream end of *comEB* should not be translated. In contrast, in the *comEB^Δ84^, comEB^Δ486^* and *comEB^Δloopout^* constructs, all of which retain higher transformability than *ΔcomEB::ery,* these final 7 codons are translated beginning from the start of the *comEB* open reading frame. Thus, it appeared possible that translation through a sequence embedded near the end of *comEB* was needed to engage an RBS, which cannot work efficiently on its own. To directly test the need for translation of *comEB,* we inserted a stop codon (TAA) in *comEB,* substituting for the native GAA at codon 72. This strain *(comEB^STOP^)* had a transformation frequency 100-fold lower than the wild-type strain (Table 1). Thus, in two constructs *(ΔcomEB::ery* and *comEB^STOP^*), when translation of *comEB* was interrupted upstream of the final 7 codons, transformation was deficient, showing that translation through these codons is required for the translation of *comEC.*

It has been reported that in the absence of ComEB, the essential transformation protein ComGA does not localize near the poles of competent cells (Burghard-Schrod *et al*., 2020) where it normally tends to reside (Hahn *et al*., 2005, Kidane & Graumann, 2005). This would suggest a role for ComEB aside from its lack of an essential role in transformation *per se.* To re-examine this issue, we expressed a functional fusion of green fluorescent protein (GFP) to the C-terminus of ComGA from its native promoter, placed at the ectopic *amyE* locus (Hahn *et al*., 2005). Fig. 4 shows that in our hands, the polar and near-polar localizations of ComGA-GFP are not noticeably affected by the absence of ComEB. We have no explanation for the discrepancy between our results and the imaging data of Burghard-Schrod et al (2020).

**Fig. 4.**
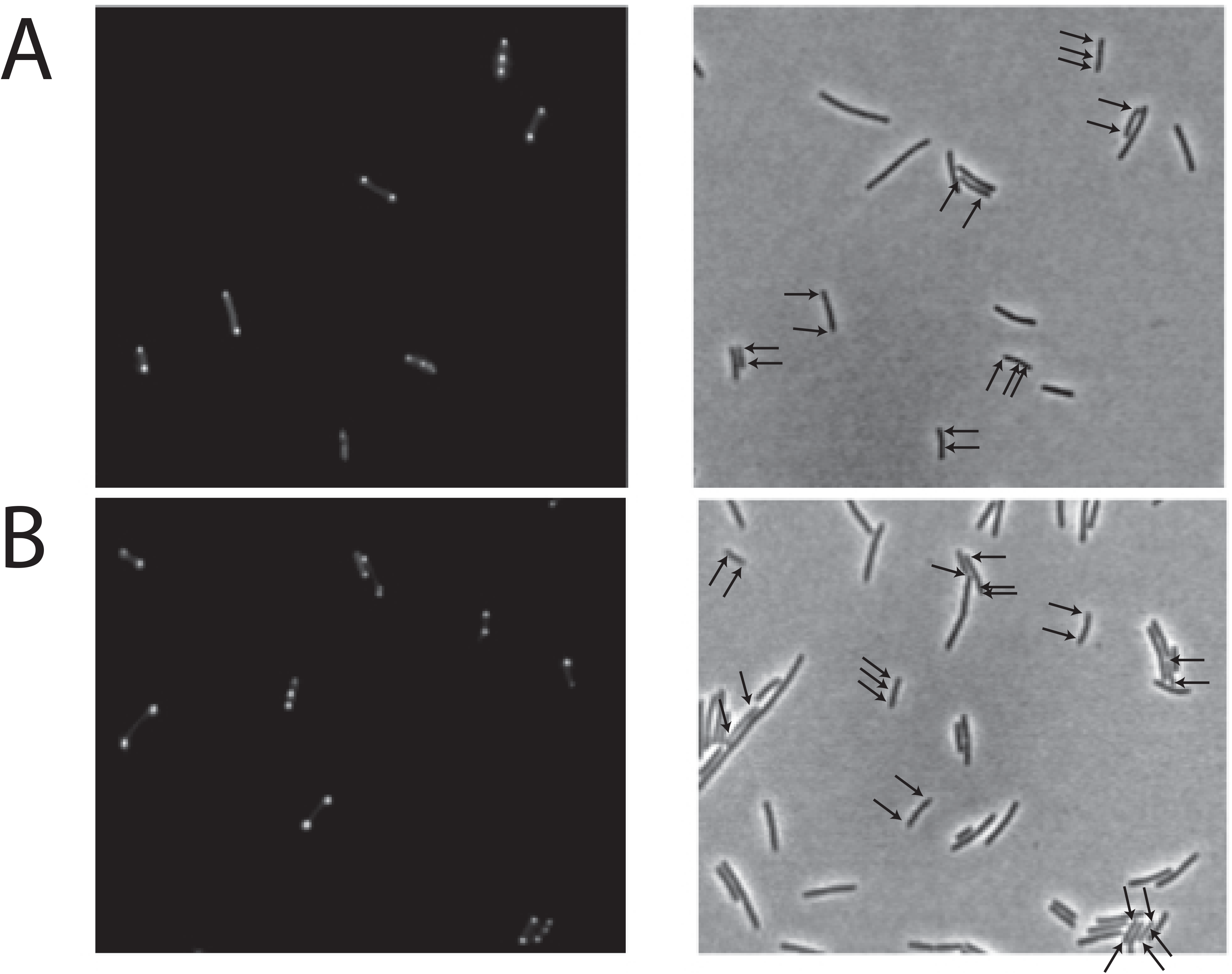
ComGA-GFP localization in a *ΔcomEB* strain. Isogenic (A) wild-type (BD2899) and (B) *ΔcomEB::ery* (BD8899) strains carrying C-terminal fusions of GFP to ComGA and transcribed by the promoter of the *comG* operon, were grown to competence and imaged for fluorescence and by phase contrast. The left and right figures in each panel show fluorescence and phase contrast images respectively. Arrows indicate the locations of the dots on the phase contrast images. As expected, a minority of the cells exhibit dots because competence is bimodally expressed.

## Discussion

A significant finding of this study is that the ComEB protein plays no detectable role in transformation. However, some interruptions of the *comEB* coding sequence markedly reduce the transformation frequency because they interfere with the synthesis of the essential ComEC protein. Promoter mapping previously revealed that a major promoter drives transcription of the entire *comE* operon and a possible minor promoter lies between *comEA* and *comEB* (Hahn *et al*., 1993), indicating that deletion of *comEB* should not interfere with the transcription of *comEC* by removing a promoter. The transformation deficiency of the *comEB^STOP^* strain further argues that the polarity effect is not exerted on the level of transcription.

Based on our results, two features of *comEB* are likely to be needed for the production of ComEC; translation of the reading frame and the presence of a sequence within the final 21 bases of *comEB* that acts as a suboptimal or inaccessible RBS for *comEC*. In fact, a sequence (ACGAGCT) is present within these final few residues of *comEB,* containing 5 out of 7 bases complementary to a sequence near the 3’ terminus of *B. subtilis* 16s rRNA (Figs. 1 and S1). Perfect complementarity (AGGAGGT), defines the canonical Shine-Dalgarno motif for *B. subtilis* (Shine & Dalgarno, 1975, Abolbaghaei *et al*., 2017). The positionw of this possible *comEB* Shine-Dalgarno motif with respect to the *comEC* start codon (D_tostart_ as defined by Prabhakaran *et al.* (2015)), is 22 residues, a few more than the typical 18-19 residues but within the permissible range. This sequence (Fig. 5), which constitutes a suboptimal RBS, is present in the three constructs that exhibit normal or near normal transformation (*comEB^Δ84^, comEB^Δ486^* and *comEB^Δloopout^*).

**Fig. 5.**
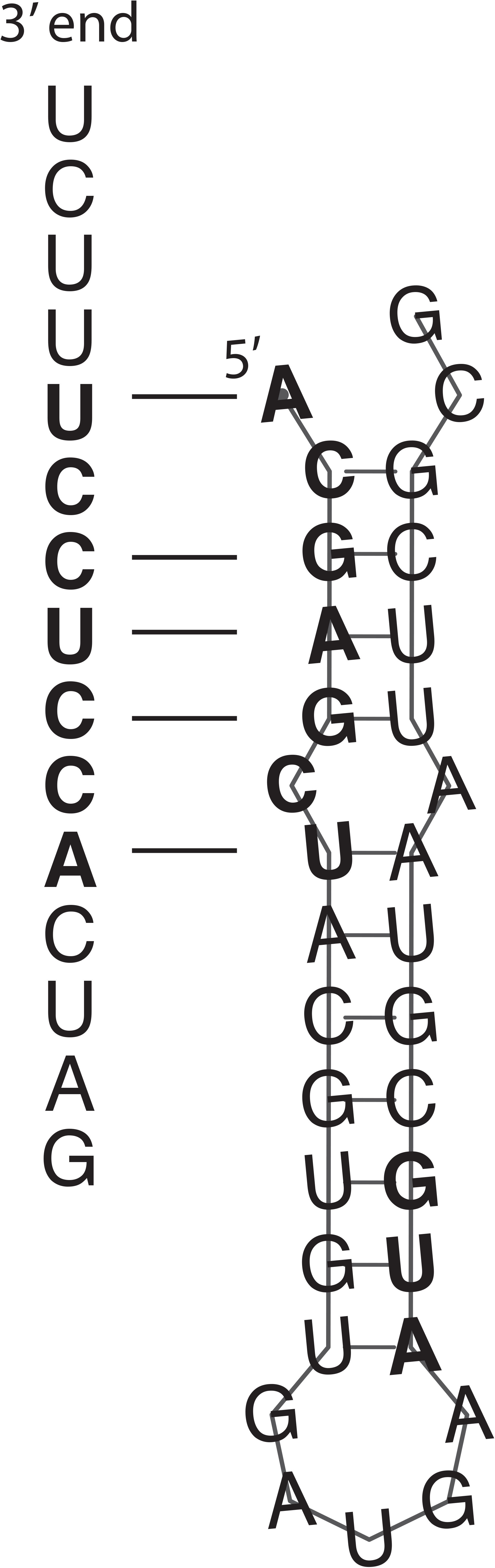
The final 12 residues of the *comEB*_reading frame, the intergenic 6 residues and the first 14 residues of *comEC* are shown as mRNA. The proposed ACGAGCU Shine-Dalgarno sequence and the *comEC* start codon are shown in boldface. The structure was predicted using RNAfold (Lorenz *et al*., 2011). Also shown are the 15 most 3-terminal residues of the *B. subtilis* 16s rRNA, with proposed complementary residues in boldface (Abolbaghaei *et al*., 2017). The D_tostart_ distance defined by (Prabhakaran *et al*., 2015) is determined by counting from the 3’ terminus of rRNA to the start codon of the relevant gene. In this case the distance is 22 residues.

Although the deficient phenotypes of the *comEB^STOP^* and *ΔcomEB::ery* strains show that translation of *comEB* is important for the expression of *comEC,* the fact that they are significantly more transformable than a *comEC* deletion strain (Table 1), argues that ribosomes can probably load inefficiently without upstream translation at an RBS within *comEB,* probably the ACGAGCT sequence noted above. In fact the *ΔcomEB::tet* construct used by Burghard-Schrod et al (2020) removed this putative RBS and retained no residual transformation, in support of this idea. We have consistently observed that the *comEB^Δ486^* strain exhibits a higher transformation frequency than the wild type strain, whereas that of the *comEB^Δloopout^* strain is reduced several fold (Table 1). These variations are consistent with the finding that translational coupling is affected by local sequence in subtle and complex ways (Levin-Karp *et al*., 2013, Buchan & Stansfield, 2007).

A translating ribosome will terminate translation from the *comEB* transcript at the sequence **UGA UGA** *AUG.* After release of the nascent ComEB protein, a ribosome may be favorably positioned to engage the RBS centered at the suboptimal ACGAGCT motif to initiate the translation of *comEC* either by direct interaction or because the local concentration of ribosomes is elevated following their release. It is also possible that ribosomes unfold an RNA structure during upstream translation, exposing sequences for *comEC* translation (Rex *et al*., 1994, Spanjaard & van Duin, 1989, Huber *et al*., 2019, Saito *et al*., 2020). In fact, analysis of the downstream sequence of the *comEB* open reading frame using RNAfold (Lorenz *et al*., 2011), suggests a structure, with a calculated free energy of −10 kcal/mol and base pairing that would sequester the putative SD motif and the *comEC* start codon (Fig. 5). Of course, these possibilities are not mutually exclusive; both the non-canonical sequence of the RBS and the presence of the stem-loop may be overcome as a result of *comEB* translation. Whatever the precise mechanism, translational coupling of *comEB* and *comEC* renders the production of ComEC, and thus transformability, largely dependent on translation of *comEB.*

Translational coupling may serve to fine-tune the synthesis of ComEC independently of the DNA receptor ComEA, which may be needed in relatively large amounts to ensure the uptake of DNA. Examination of the *comE* operon in other bacteria suggests that coupling of *comEB* and *comEC* is widespread. *Lactobacillus plantarum* and *B. anthracis* are similar to *B. subtilis* as their *comEB* and *comEC* reading frames are nearly contiguous (Fig. 1), and so are *Leuconstoc mesenteroides, L. lactis* and *B. licheniformis* (not shown). Other firmicutes, e.g. *L. amylovorus* and *S. pneumoniae* (Fig. 1), as well as *B. pumilus, S. mutans, Geobacillus thermoleovorans* and the Gram negative *Thermus thermophilus* (not shown) have actual overlaps between the *comEB* and *comEC* coding sequences. *Staphylococcus aureus* has a well-spaced arrangement of the genes, but with an apparently weak Shine-Dalgarno sequence between *comEB* and *comEC.* It seems that a variety of mechanisms have evolved to down-regulate the synthesis of ComEC relative to that of ComEA.

Interestingly, the *B. subtilis comFC* gene is preceded by *comFB* that encodes a protein of unknown function dispensable for transformation (Sysoeva *et al*., 2015). As pointed out previously (Sysoeva *et al*., 2015), *comFB* and *comFC* overlap, suggesting that they are also translationally coupled and this arrangement may serve to adjust the expression of ComFC with respect to ComFA, both of which are essential for transformation. Also, in the *comG* operon, which encodes the transformation pilus, the *comGC, comGD, comGF* and *comGG* genes successively overlap, as noted previously (Albano *et al*., 1989). Thus in *B. subtilis,* all of the major operons required for transformation *(comG, comF* and *comE),* exhibit the potential for translational coupling that may serve to fine-tune expression stoichiometries for proteins that work in supra-molecular complexes, partially detaching the translation of downstream genes from strict transcriptional control.

These data establish the dispensability of ComEB for transformation, but beg the question of a possible secondary role for ComEB in the context of competence; why do many firmicutes encode ComEB in the ComE operon, placing the transcription of *comEB* under competence control? It was suggested that ComEB aids in the proper localization of ComGA in the competent cell, but our results do not support that conclusion. Perhaps ComEB, which is a dCMP deaminase (Burghard-Schrod *et al*., 2020, Oehlenschlaeger *et al*., 2015), plays a role in scavenging pyrimidines, including those released by degradation of the non-transforming strand during DNA transport. In fact, the products of this degradation include 5’ deoxyribomononucleotides, released into the medium (Dubnau & Cirigliano, 1972). In this regard it is interesting that ComFC, a conserved protein that is required for DNA uptake, contains a domain that resembles a type 1 phosphoribosyl transferase (L. Celma, S. Marsin, P. Radicella and S. Quevillon-Cheruel, personal communication), another enzyme that plays a role in nucleic acid base salvage. Also, transcriptional profiling suggests that the genes encoding xanthine phosphoribosyl transferase and a xanthine transporter, *pbuX* and *xpt* respectively, are induced by ComK, and thus expressed in the competent state (Berka *et al*., 2002). These various scavenging enzymes may increase fitness in the context of competence, particularly under conditions of nutritional scarcity, apart from any direct role they may or may not play in DNA transport.

These speculations are relevant to the recurring debate over the role(s) of transformation; is it for food or for genetic information or for both, as suggested by Redfield (1993)? Perhaps in at least some bacteria, an enhanced ability to use the nontransforming donor DNA and the recipient DNA displaced by integration for nutrition, accompanies the induction of competence for the acquisition of genetic information, a parsimonious use of resources to enhance fitness.

## Experimental Procedures

### Bacteria, growth and transformation

All of the strains (Table 2) described in this report are derivatives of *B. subtilis* 168 and are isogenic with IS75 (*his leu met).* For growth purposes other than transformation, the bacteria were grown in LB medium (Kearns & Losick, 2005). Growth to competence was by the “two-step” procedure as described in Dubnau and Davidoff-Abelson (1971) except that fresh, unfrozen cells were used for transformation. After growth to competence, 1 ml of culture was incubated with 1 μg of chromosomal DNA from BD170 (*thr trp)* for 30 minutes at 37° C. Transformants were selected on glucose minimal medium agar lacking leucine for selection of Leu^+^ prototrophy. Transformation frequencies were calculated as the number of prototrophic transformants ml^-1^ divided by the number of colony forming units ml^-1^.

**Table 2.**
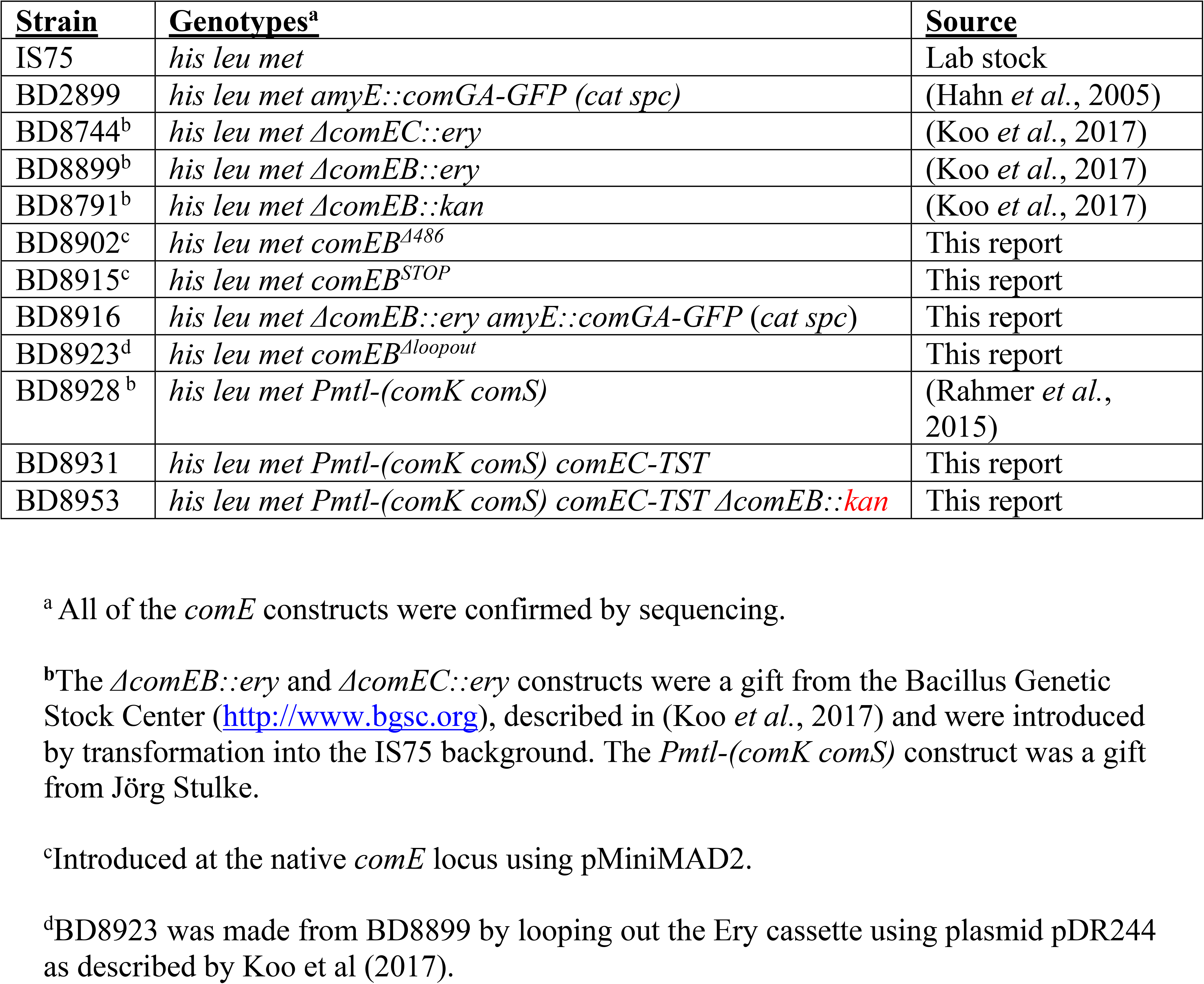
Strains

Strains
| **Strain** | **Genotypes<sup>a</sup>** | **Source** |
| --- | --- | --- |
| IS75 | *his leu met* | Lab stock |
| BD2899 | *his leu met amyE::comGA-GFP (cat spc)* | (Hahn *et al.*, 2005) |
| BD8744<sup>b</sup> | *his leu met ΔcomEC::ery* | (Koo *et al.*, 2017) |
| BD8899<sup>b</sup> | *his leu met ΔcomEB::ery* | (Koo *et al.*, 2017) |
| BD8791<sup>b</sup> | *his leu met ΔcomEB::kan* | (Koo *et al.*, 2017) |
| BD8902<sup>c</sup> | *his leu met comEB<sup>Δ486</sup>* | This report |
| BD8915<sup>c</sup> | *his leu met comEB<sup>STOP</sup>* | This report |
| BD8916 | *his leu met ΔcomEB::ery amyE::comGA-GFP (cat spc)* | This report |
| BD8923<sup>d</sup> | *his leu met comEB<sup>Δloopout</sup>* | This report |
| BD8928<sup>b</sup> | *his leu met Pmtl-(comK comS)* | (Rahmer *et al.*, 2015) |
| BD8931 | *his leu met Pmtl-(comK comS) comEC-TST* | This report |
| BD8953 | *his leu met Pmtl-(comK comS) comEC-TST ΔcomEB::kan* | This report |

<sup>a</sup> All of the *comE* constructs were confirmed by sequencing.
<sup>b</sup> The *ΔcomEB::ery* and *ΔcomEC::ery* constructs were a gift from the Bacillus Genetic Stock Center (http://www.bgsc.org), described in (Koo *et al.*, 2017) and were introduced by transformation into the IS75 background. The *Pmtl-(comK comS)* construct was a gift from Jörg Stulke.
<sup>c</sup> Introduced at the native *comE* locus using pMiniMAD2.
<sup>d</sup> BD8923 was made from BD8899 by looping out the Ery cassette using plasmid pDR244 as described by Koo *et al.* (2017).

### Introduction of markerless constructs to the *B. subtilis* chromosome

Mutant genes of interest were cloned into pMiniMAD2, a gift from Dan Kearns (Indiana University). After transformation with selection for erythromycin (5 μg/ml), markerless mutants were isolated essentially as described by Patrick and Kearns (2008). In all cases the mutant constructs were confirmed by sequencing.

### Mutant constructs

The primers used for the construction of mutant plasmids are described in Table 3. All constructs were verified by sequencing.

**Table 3.**
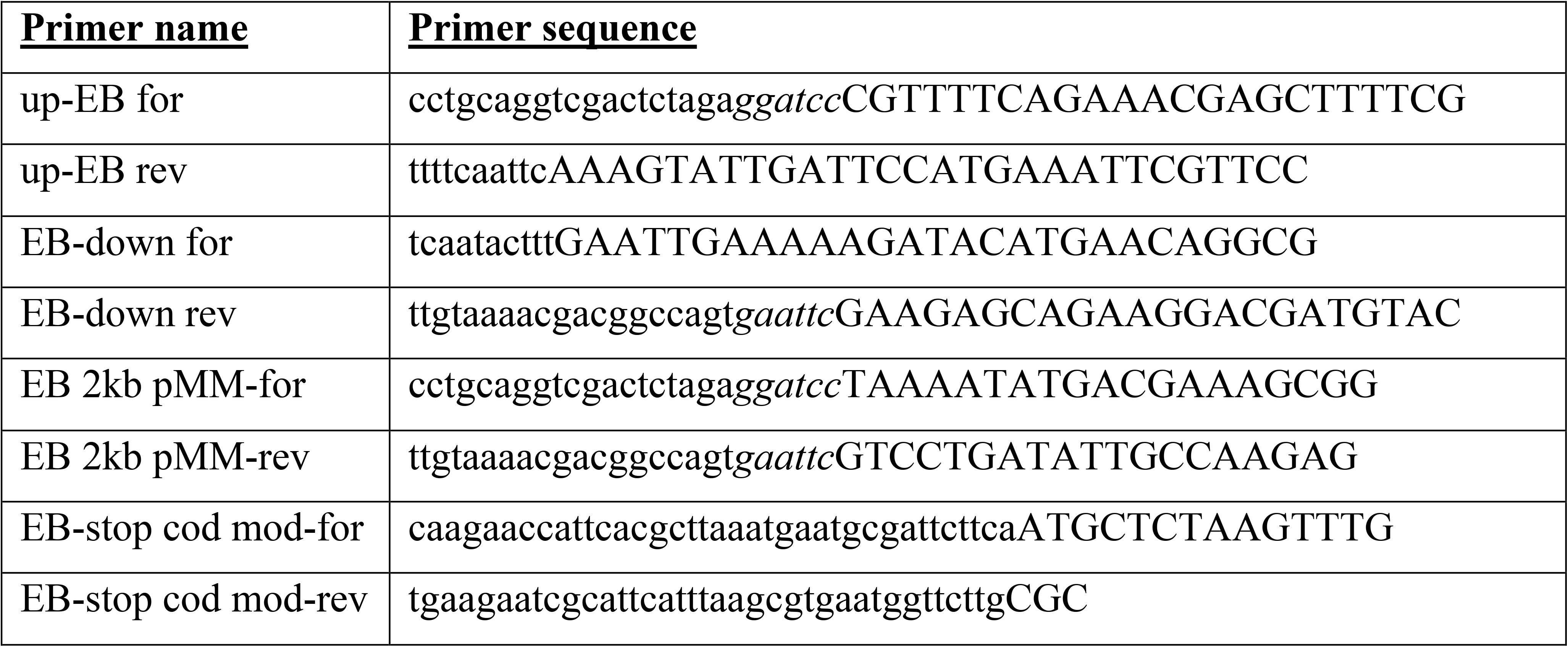
Primers

***comEB^Δloopout^* (BD8923)** was constructed from BD8899 (Table 2) by looping out the erythromycin resistance cassette using plasmid pDR244, as described by Koo et al (2017).

***comEB^Δ486^* (BD8902)** was obtained by PCR amplification of the 1 kb regions upstream and downstream of the 486 *comEB* reidues to delete by using respectively the primer pairs up-EBfor/up-EBrev and EB-downfor/EB-downrev. The two resulting fragments were cloned using the NEBuilder HiFi DNA assembly kit into pMiniMad2 plasmid previously digested using the restriction sites *BamHI* and *Eco*RI. The resulting plasmid pED2386 was moved into the chromosome of *B. subtilis* at the native locus as described previously (Mukherjee *et al.*, 2013).

***comEB^STOP^* (BD8915)** was constructed by PCR amplification of the 1 kb regions upstream and downstream of the *comEB* glutamate amino acid at position 72 by using the primer pair EB 2kb pMM-for and EB 2kb MM-rev. The resulting fragment was cloned using the NEBuilder HiFi DNA assembly kit (NEW ENGLAND BioLabs) into the pMiniMAD2 plasmid previously digested using the restriction sites *Bam*HI and *Eco*RI. Mutagenesis was carried out using the Phusion site directed mutagenesis kit (Thermo Fisher) and primer pair EB-stop cod mod-for and EB stop-cod mod-rev. The resulting plasmid pED2392 was moved into the chromosome of *B. subtilis* at the native locus as above.

### ComEC-TST fusion

The fusion construct comEC-Twin strep tag (plasmid pED2385) was generated from a synthesized fragment purchased from Genscript USA that contained two strep tag II sequences, separated by a linker and flanked with *BamHI* and *SacI* restriction sites. A 998 bp DNA fragment carrying the C terminus of *comEC* was cloned into the pUC19 plasmid using the restriction sites *Sal*I and *Bam*HI. The synthetic twin strep tag fragment (93 bp) was added to the *comEC* gene in the resulting plasmid with the restriction sites *BamHI* and *SacI.* The entire fragment carrying the *comEC*-twin strep tag fusion construct was then inserted into the *SalI* and *Sac*I sites of pUC-CM (pED2384) previously digested using the same restriction sites. The resulting plasmid pED2389, was then transformed into BD8928 with selection for chloramphenicol-resistance by Campbell-like transformation, placing the fusion construct under control of the native *comEC* promoter to produce BD8931. The *ΔcomEB::kan* (BD8791) construct was in turn introduced by transformation to form BD8953.

### Microscopy

After growth to the competent state, 1 μl of culture was deposited on an agarose pad and imaged by phase microscopy as well as for GFP fluorescence, using a Nikon T*i* microscope. The images in Fig. 3 were collected using an Orca Flash 4.0 camera (Hamamatsu) and processed identically using Elements software (Nikon) and Photoshop (Adobe).

### Western blotting

We detected ComEC using an antiserum (IBA Lifesciences) against a C-terminal twin-strep tag fusion to ComEC in the *Pmtl-(comK comS)* background (BD8931) and in an isogenic *comEB::ery* strain (BD8953). Membrane fractions were isolated as described in Draskovic et al (2005), except that the membrane fractions were simply pelleted by centrifugation at 100,000 x g and not purified further. In addition to the isogenic *comEB^+^*strain, a *Pmtl-(comK comS)* strain with no tag (BD8928) was included to confirm the identification of the ComEC-TST signal. The strains were grown in LB-broth to mid-log and induced by the addition of mannitol (1%). After 90 minutes the cells were harvested for Western blotting.

## Acknowledgements

This work was supported by NIH grant R01GM057720. We thank Mathew Neiditch and the members of the Dubnau lab for helpful comments and discussion. We thank Daniel Ziegler at the Bacillus Genetic Stock Center and Jörg Stulke for generously providing strains. The authors have no conflicts of interest to declare that are relevant to the content of this article.

## Author contributions

M. D. S. and J. H. performed all of the experimental work in this work. D. D. supervised the work and all the authors contributed to planning experiments, analyzing data and to writing the paper.

## References

Abolbaghaei, A., J. R. Silke & X. Xia, (2017) How Changes in Anti-SD Sequences Would Affect SD Sequences in Escherichia coli and Bacillus subtilis. G3 (Bethesda) 7: 1607–1615.

Albano, M., R. Breitling & D. A. Dubnau, (1989) Nucleotide sequence and genetic organization of the *Bacillus subtilis* comG operon. J Bacteriol 171: 5386–5404.

Berka, R. M., J. Hahn, M. Albano, I. Draskovic, M. Persuh, X. Cui, A. Sloma, W. Widner & D. Dubnau, (2002) Microarray analysis of the *Bacillus subtilis* K-state: genome-wide expression changes dependent on ComK. Mol Microbiol 43: 1331–1345.

Blokesch, M., (2016) Natural competence for transformation. Curr Biol 26: R1126–R1130.

Buchan, J. R. & I. Stansfield, (2007) Halting a cellular production line: responses to ribosomal pausing during translation. Biol Cell 99: 475–487.

Burghard-Schrod, M., S. Altenburger & P. L. Graumann, (2020) The *Bacillus subtilis* dCMP deaminase ComEB acts as a dynamic polar localization factor for ComGA within the competence machinery. Mol Microbiol 113: 906–922.

Draskovic, I. & D. Dubnau, (2005) Biogenesis of a putative channel protein, ComEC, required for DNA uptake: membrane topology, oligomerization and formation of disulphide bonds. Mol Microbiol 55: 881–896.

Dubnau, D. & M. Blokesch, (2019) Mechanisms of DNA Uptake by Naturally Competent Bacteria. Annu Rev Genet 53: 217–237.

Dubnau, D. & C. Cirigliano, (1972) Fate of transforming DNA following uptake by competent *Bacillus subtilis.* III. Formation and properties of products isolated from transformed cells which are derived entirely from donor DNA. J Mol Biol 64: 9–29.

Dubnau, D. & R. Davidoff-Abelson, (1971) Fate of transforming DNA following uptake by competent *Bacillus subtilis.* I. Formation and properties of the donor-recipient complex. J Mol Biol 56: 209–221.

Gangel, H., C. Hepp, S. Muller, E. R. Oldewurtel, F. E. Aas, M. Koomey & B. Maier, (2014) Concerted spatio-temporal dynamics of imported DNA and ComE DNA uptake protein during gonococcal transformation. PLoS Pathog 10: e1004043.

Hahn, J., G. Inamine, Y. Kozlov & D. Dubnau, (1993) Characterization of *comE,* a late competence operon of *Bacillus subtilis* required for the binding and uptake of transforming DNA. Mol Microbiol 10: 99–111.

Hahn, J., B. Maier, B. J. Haijema, M. Sheetz & D. Dubnau, (2005) Transformation proteins and DNA uptake localize to the cell poles in *Bacillus subtilis*. Cell 122: 59–71.

Huber, M., G. Faure, S. Laass, E. Kolbe, K. Seitz, C. Wehrheim, Y. I. Wolf, E. V. Koonin & J. Soppa, (2019) Translational coupling via termination-reinitiation in archaea and bacteria. Nat Commun 10: 4006.

Inamine, G. S. & D. Dubnau, (1995) ComEA, a *Bacillus subtilis* integral membrane protein required for genetic transformation, is needed for both DNA binding and transport. J Bacteriol 177: 3045–3051.

Kearns, D. B. & R. Losick, (2005) Cell population heterogeneity during growth of *Bacillus subtilis*. Genes Dev 19: 3083–3094.

Kidane, D. & P. L. Graumann, (2005) Intracellular protein and DNA dynamics in competent *Bacillus subtilis* cells. Cell 122: 73–84.

Koo, B. M., G. Kritikos, J. D. Farelli, H. Todor, K. Tong, H. Kimsey, I. Wapinski, M. Galardini, A. Cabal, J. M. Peters, A. B. Hachmann, D. Z. Rudner, K. N. Allen, A. Typas & C. A. Gross, (2017) Construction and Analysis of Two Genome-Scale Deletion Libraries for Bacillus subtilis. Cell Syst 4: 291–305 e297.

Levin-Karp, A., U. Barenholz, T. Bareia, M. Dayagi, L. Zelcbuch, N. Antonovsky, E. Noor & R. Milo, (2013) Quantifying translational coupling in E. coli synthetic operons using RBS modulation and fluorescent reporters. ACS Synth Biol 2: 327–336.

Lorenz, R., S. H. Bernhart, C. Honer Zu Siederdissen, H. Tafer, C. Flamm, P. F. Stadler & I. L. Hofacker, (2011) ViennaRNA Package 2.0. Algorithms Mol Biol 6: 26.

Maier, B., (2020) Competence and Transformation in *Bacillus subtilis*. Curr Issues Mol Biol 37: 57–76.

Matthey, N. & M. Blokesch, (2016) The DNA-Uptake Process of Naturally Competent *Vibrio cholerae*. Trends Microbiol 24: 98–110.

Mukherjee, S., P. Babitzke & D. B. Kearns, (2013) FliW and FliS function independently to control cytoplasmic flagellin levels in *Bacillus subtilis*. J Bacteriol 195: 297–306.

Oehlenschlaeger, C. B., M. N. Lovgreen, E. Reinauer, E. Lehtinen, M. L. Pind, P. Harris, J. Martinussen & M. Willemoes, (2015) *Bacillus halodurans* Strain C125 Encodes and Synthesizes Enzymes from Both Known Pathways To Form dUMP Directly from Cytosine Deoxyribonucleotides. Appl Environ Microbiol 81: 3395–3404.

Patrick, J. E. & D. B. Kearns, (2008) MinJ (YvjD) is a topological determinant of cell division in *Bacillus subtilis*. Mol Microbiol 70: 1166–1179.

Prabhakaran, R., S. Chithambaram & X. Xia, (2015) Escherichia coli and Staphylococcus phages: effect of translation initiation efficiency on differential codon adaptation mediated by virulent and temperate lifestyles. J Gen Virol 96: 1169–1179.

Provvedi, R. & D. Dubnau, (1999) ComEA is a DNA receptor for transformation of competent *Bacillus subtilis*. Mol Microbiol 31: 271–280.

Rahmer, R., K. Morabbi Heravi & J. Altenbuchner, (2015) Construction of a Super-Competent Bacillus subtilis 168 Using the P mtlA -comKS Inducible Cassette. Frontiers in microbiology 6: 1431.

Redfield, R. J., (1993) Genes for breakfast: the have-your-cake-and-eat-it-too of bacterial transformation. J Hered 84: 400–404.

Rex, G., B. Surin, G. Besse, B. Schneppe & J. E. McCarthy, (1994) The mechanism of translational coupling in *Escherichia coli.* Higher order structure in the *atpHA* mRNA acts as a conformational switch regulating the access of de novo initiating ribosomes. J Biol Chem 269: 18118–18127.

Saito, K., R. Green & A. R. Buskirk, (2020) Ribosome recycling is not critical for translational coupling in Escherichia coli. Elife 9.

Schwarzenlander, C., W. Haase & B. Averhoff, (2009) The role of single subunits of the DNA transport machinery of *Thermus thermophilus* HB27 in DNA binding and transport. Environ Microbiol 11: 801–808.

Shine, J. & L. Dalgarno, (1975) Terminal-sequence analysis of bacterial ribosomal RNA. Correlation between the 3’-terminal-polypyrimidine sequence of 16-S RNA and translational specificity of the ribosome. Eur J Biochem 57: 221–230.

Spanjaard, R. A. & J. van Duin, (1989) Translational reinitiation in the presence and absence of a Shine and Dalgarno sequence. Nucleic Acids Res 17: 5501–5507.

Sysoeva, T. A., L. B. Bane, D. Y. Xiao, B. Bose, S. S. Chilton, R. Gaudet & B. M. Burton, (2015) Structural characterization of the late competence protein ComFB from *Bacillus subtilis*. Biosci Rep 35.

